# Somatostatin Receptors Shape Insulin and Glucagon Output within the Pancreatic Islet through Direct and Paracrine Effects

**DOI:** 10.1101/2025.11.13.688371

**Authors:** Ryan G. Hart, Jordan J. Lee, Karen Zhai, Sharlene Lee, Rashita Chauhan, Aidean Hosseini, Austin D. Nguyen, Mark O. Huising

**Affiliations:** Department of Neurobiology, Physiology and Behavior, University of California Davis, Davis, California, USA; Department of Physiology and Membrane Biology, University of California Davis, Davis, California, USA

## Abstract

**Aims/Hypothesis:** Pancreatic delta cells secrete somatostatin (SST), which can inhibit both alpha and beta cells of the pancreatic islet. By controlling insulin and glucagon release, delta cells play an important role in maintaining nutrient homeostasis. However, the mechanism by which a single inhibitory hormone inhibits both alpha and beta cells, which are often considered as functional antagonists in the counterregulatory control of blood glucose, has been a physiological riddle. Here, we solve this riddle through assessment of the contributions of alpha and beta cell specific somatostatin receptors to cell intrinsic behaviors and hormone release.

**Methods:** Islets from mice constitutively expressing fluorescent sensors reporting on cyclic AMP and Ca^2+^ in both alpha and beta cells were imaged using stimuli to mimic the post-prandial state of a meal consisting of glucose and amino acids. This approach was coupled with cell specific somatostatin receptor antagonists to identify how somatostatin inhibits alpha and beta cell hormone output through modulation of cAMP and Ca^2+^ secondary messengers and paracrine interactions.

**Results:** Our results support and extend prior observation that somatostatin receptor 2 (SSTR2) is the only somatostatin receptor expressed by alpha cells, while somatostatin receptor 3 (SSTR3) is the only receptor expressed by mouse beta cells. Interestingly, SSTR2 and SSTR3 regulate downstream cAMP and Ca^2+^ signaling cascades differently within alpha and beta cells of intact islets. Stimulation of somatostatin receptors robustly inhibits cyclic AMP in alpha or beta cells. In contrast, stimulation of SSTR2 inhibits alpha cell Ca^2+^ with significantly greater potency compared to inhibition of beta cell Ca^2+^ via SSTR3. Despite the absence of SSTR2 on beta cells, blocking alpha cell SSTR2 during nutrient stimulation resulted in a significant increase in insulin release downstream of local release of glucagon.

**Conclusions/Interpretation:** Our observations address the physiological riddle of the delta cell’s role during the post-prandial phase where we demonstrate that somatostatin primarily inhibits alpha cell cAMP and Ca^2+^ via SSTR2, preventing glucagon release. Blocking SSTR2 resulted in an increase in locally released glucagon, which coupled with muted ability for SSTR3 to inhibit beta cell calcium under strong nutrient stimulation, results in potentiation of glucose stimulated insulin secretion from the beta cell. We conclude that the role of delta cells under nutrient stimulation is to modulate the volume of insulin release by tuning the strength of intra-islet paracrine potentiation of insulin secretion by glucagon, mediated via beta cell GLP1R.

**Research in Context:** *What is already known about this subject?:*

- Somatostatin released by delta cells can attenuate glucagon release from the alpha cells and insulin release from the beta cells of the islet through inhibitory somatostatin receptors.
- Somatostatin receptor 2 is a major receptor expressed on the mouse alpha cell surface, while mouse beta cells express SSTR3 on their primary cilia.
- Paracrine signaling by alpha cell glucagon potentiates insulin release through the engagement of stimulatory glucagon-like peptide 1 receptors on beta cells.

*What are the key questions?:*

- How do the cell specific somatostatin receptors of the alpha cell (SSTR2) and beta cell (SSTR3) differentially regulate cell-intrinsic Ca2+ and cAMP signaling and the indirect paracrine pathways that shape insulin secretion in the post-prandial state?

*What are the new findings?:*

- Alpha cells of the mouse islet express exclusively SSTR2 and not SSTR3.
- Somatostatin is a more potent inhibitor of cytosolic calcium activity via alpha cell SSTR2 compared to beta cell SSTR3.
- Preventing somatostatin inhibition of the alpha cell in the presence of elevated glucose and amino acids potentiates insulin release through the engagement of glucagon-like peptide 1 receptors on the beta cell by glucagon.

*How might this impact on clinical practice in the foreseeable future?:*

- Our findings indicate that local feedback inhibition of beta cells is accomplished by tuning the strength of the paracrine potentiation of locally released glucagon. This is relevant in the context of developing treatments for hypoglycemia with alpha cell specific SSTR2-specific antagonists that could potentiate insulin release and potentially exacerbate hypoglycemic episodes.

**Graphical Abstract:** 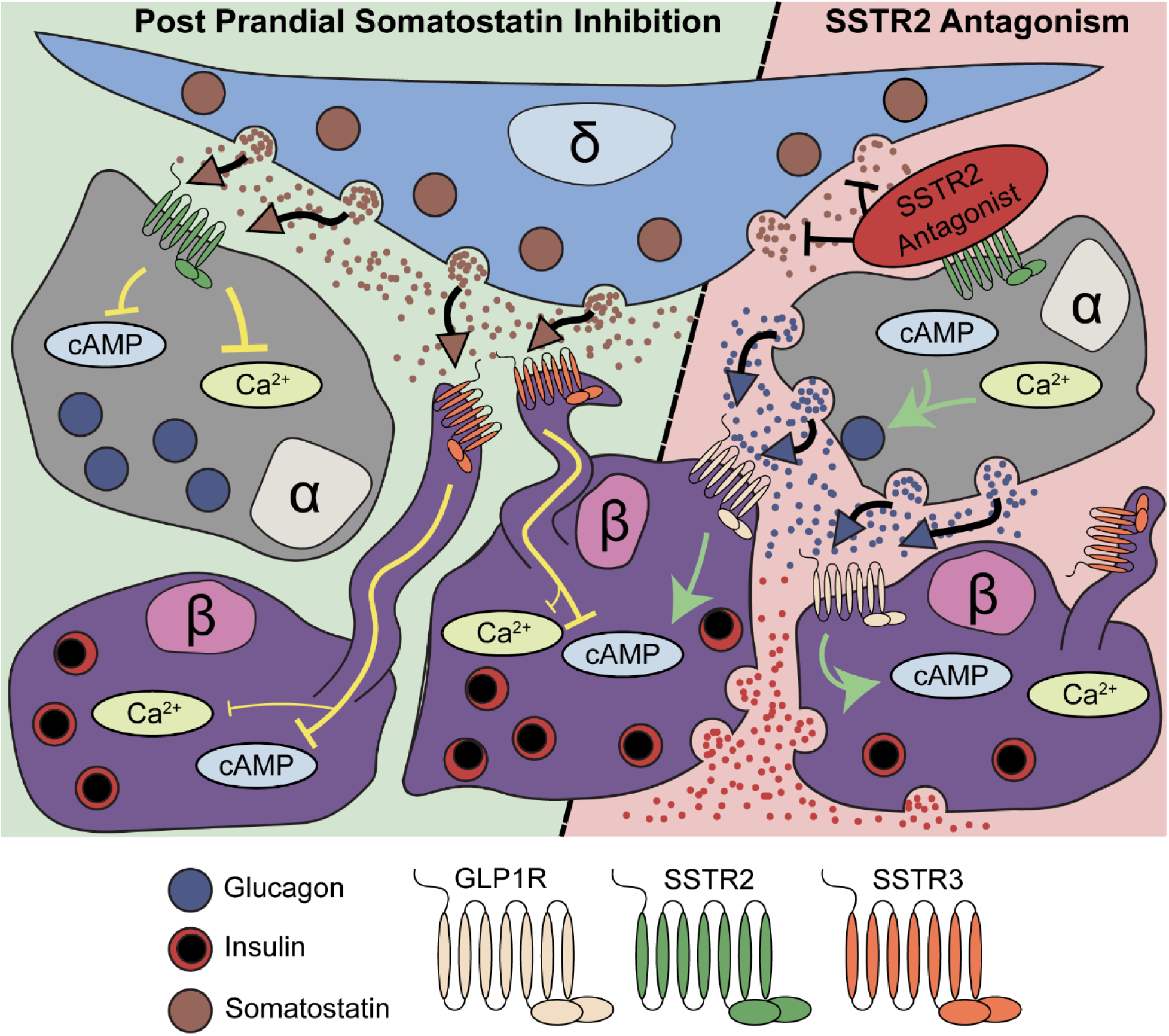

## Introduction

Within the pancreatic islets of Langerhans, a tightly regulated network of cells is responsible for maintaining nutrient homeostasis [1]. Two of these cell types, the alpha (α) and beta (β) cells, are considered to achieve this by balancing the secretion of glucagon and insulin respectively [2]. Glucagon is an important counterregulatory hormone that elevates blood glucose under hypoglycemic conditions by stimulating hepatic glucose production [3]. Insulin promotes the anabolism of digested and absorbed nutrients to facilitate clearance and lowering of circulating blood glucose [4]. The delta (δ) cell is the third main islet endocrine cell type, secreting the hormone somatostatin (SST) [5], which was recognized as a local inhibitor of α and β cell hormone secretion shortly after its discovery as the inhibitor of growth hormone release [6]. Recently we demonstrated how the feedback inhibition provided by δ cells is coordinated with β cells [7] and sets the homeostatic set point for blood glucose between meals [8]. However, how a single inhibitory hormone (SST) directs feedback inhibition to both α and β cells simultaneously, has not been satisfactorily explained. Under a counterregulatory model where glucagon and insulin are released at low and high glucose concentrations, respectively, it would not make physiological sense to contemporaneously inhibit secretion of both hormones. However, a series of recent publications [9–13] build on long-standing observations that glucagon paradoxically stimulates insulin secretion [14] and make a compelling case that glucagon during the fed state serves as an important local amplifier of nutrient-stimulated insulin secretion via the activation of the predominantly β cell Glucagon Like Peptide-1 Receptor (GLP1R) [14]. These observations necessitate a revisiting of the role of α cells as local cooperators with β cells to potentiate glucose-stimulated insulin secretion (GSIS) during the post-prandial phase [9, 10]. This model actually greatly simplifies the role of δ cell SST during the prandial phase, as inhibition of β cells and α cells would both result in the inhibition of insulin release through either direct inhibition of β cells or the inhibition of glucagon-mediated paracrine potentiation of GSIS.

Somatostatin signals through a family of 5 different inhibitory G protein coupled (Gαi GPCR) SST receptors (SSTR1-5) [15]. Comprehensive transcriptomes of FACS-purified α and β cells have established *Sstr3* as the only SSTR mRNA present in mouse β cells, whereas *Sstr2* and *Sstr3* mRNA is expressed by mouse α cells [16–19]. All SSTRs are Gαi-coupled GPCRs that are expressed on the cell membrane, except for SSTR3, which is expressed on the primary cilia [20]. Whether the marked differences in localization of SSTR2 and SSTR3 affect downstream signaling and their potency to inhibit islet hormone release has not been determined. SST-mediated activation of SSTRs will release the alpha, beta, and gamma trimeric proteins. The Gαi subunit inhibits adenylyl cyclase and reduces the production of the secondary messenger cyclic adenosine monophosphate (cAMP) [21]. Somatostatin is capable of inhibiting cytosolic calcium (Ca^2+^) via the activation of inward rectifying potassium (GIRK) channels through beta/gamma trimeric protein signaling events in some cell types [22], although recent observations indicate that Sodium Potassium ATPase transporters are the more important mechanism by which SST inhibits intracellular Ca^2+^ in primary β cells [23].

Several gaps in our understanding of SST-mediated feedback remain, including if SST inhibits insulin secretion through the direct inhibition of β cells via SSTR3 or indirectly by attenuation of the paracrine stimulation of insulin release by glucagon from α cells. We also do not know if the relative importance of inhibition by SST of intracellular Ca^2+^ and cAMP in α and β cells. To close these gaps in our understanding of the mechanisms by which SST attenuates insulin secretion, we leveraged mice expressing genetically encoded sensors reporting on both Ca^2+^ and cAMP to quantify the contributions of individual SSTRs expressed by primary α and β cells. Our findings demonstrate that SST has a major inhibitory influence on insulin release that is achieved primarily indirectly by SSTR2-mediated inhibition of α cell cAMP, Ca^2+^, and glucagon release to prevent its paracrine potentiation of insulin secretion. Somatostatin acting on β cell cilia SSTR3 also directly inhibits β cell cAMP (in part activated by local glucagon), but is less effective in inhibiting β cell Ca^2+^. These observations demonstrate that the inhibition of insulin secretion during the prandial phase is mediated by the attenuation of α cell paracrine potentiation of β cells.

## Methods

### Mouse Strains

C57BL/6N mice were group housed (up to four mice per cage) in a specific pathogen free facility on a 12-h light–dark cycle, an ambient temperature of 20–26 °C and humidity of 30–70%. Water and standard rodent chow were provided *ad libitum*. Generation of whole body expressing GCaMP6s Ca^2+^ indicator mice was accomplished by crossing heterozygous lox-stop-lox GCaMP6s mice (B6;129S6-*Gt(ROSA)26Sor*^tm96(CAG-GCaMP6s)Hze^/J, Jax 24106) [24] with a heterozygous beta-actin-Cre mouse (B6N.FVB-Tmem1636^TG(ACTB-Cre)2Mrt^/CjDswJ, Jax 019099) [25]. Once dual GCaMP6s and beta-actin-Cre mice were generated, the lox-stop-lox cassette was permanently deleted in any cell expressing beta actin, including the germ line. As a result, the functional GCaMP6s gene was then passed down to the progeny of any beta-actin-Cre x lox-Stop-lox GCaMP6s transgenic animal. Generation of whole body knock in cAMP sensor expressing mice was generated in the same manner by crossing beta-actin-Cre mice to heterozygous lox-stop-lox CAMPER mice (Gt(ROSA)26Sor^tm1(CAG-ECFP*/Rapgef3/Venus*)Kama^/J, JAX stock #032205) [26]. Islet Cell specific membrane tomato lox-stop-lox membrane GFP (mT/mG) mice were generated by crossing a heterozygous mT/mG mouse (B6.129(Cg)-Gt(ROSA)26Sor ^tm4(ACTB-tdTomato,-EGFP)Luo^/J, Jax 007676) [27] with a heterozygous mouse expressing α cell-specific (GCG-CreEr *Gc^gem1(cre/ERT2)^,* MMRRC stock #42277) [28], β cell-specific (Tg(Ucn3-Cre)KF43Gsat/Mmucd, MMRRC stock #032078) [29], or δ cell-specific (SST-Cre (*Sst^t^*^m2.1(Cre)Zjh^/J, Jax 013044) [30] Cre driver. Imaging experiments were performed with mice between 3 and 6 months of age. All mouse experiments were approved by the UC Davis Institutional Animals Care and Use Committee and were performed in compliance with the Animal Welfare Act and the Institute for Laboratory Animal Research Guide to the Care and Use of Laboratory Animals.

### Islet isolation and dissociation

Islets were isolated as described previously [7, 8]. After isolation, intact islets recovered in RPMI +10% Pen/Strep +5.5 mM glucose (complete) for 16-24 hours. For dissociation, islets were handpicked into 15 mL conical vials, gravity sedimented, and the supernatant removed. solution was added, with immediate gentle resuspension mixing 3 times with a pipette set to 200 μL and incubated at 37 degrees Celsius for 45 seconds. Islets were gently pipetted up and down 20 times using the same pipette set to 200 μl to triturate the islets. The dissociation was inactivated with 13 mL of RPMI complete and cells pelleted for 5 minutes at 500g. Dissociated cells were immediately seeded into imaging chambers.

### Static hormone-secretion assays

After overnight culture in a 5.5 mM glucose RPMI complete media. pooled islets isolated from a cohort of age-matched commercial wild type mice were picked twice into Krebs Ringer Buffer (20 mM Hepes pH 7.4, 1.2 mM KH_2_PO_4_, 25 mM NaHCO_3_, 130 mM NaCl, 5 mM KCl, 1.2 mM MgCl_2_, 1.2 mM CaCl_2_) containing 0.1% BSA and 5.5 mM glucose, then incubated at 37°C for 1 hour. For glucagon and insulin secretion, measurements were collected from the same lysates, 20 islets were picked into replicate 24-wells plate for at least 5 replicates per condition. The different stimuli were added to final concentration after the islets had been placed into the wells. The amount of insulin or glucagon present was measured using the Lumit luminescence assay kit (Promega).

### Live Imaging of Secondary Messenger Behavior

Intact islets or dissociated islet cells constitutively expressing CAMPER or GCaMP6s were set down into custom microperfusion chambers bonded to a 35 mm glass-bottom dish (Mattek, Catalog # P35G-1.5-14-C) and allowed to adhere overnight. Dissociated cultures were directly seeded into the imaging chamber 24-48 hours before imaging. Continuous perfusion of Krebs Ringer Buffer at a rate of 200 μL per minute was maintained using a customized Elveflow microfluidics system [8]. The Ca^2+^ and cAMP responses of islets over time was imaged using a Nikon Eclipse Ti2 using a 60x lens with oil for intact, and a 20x objective for dissociated cultures. Stimulation with 100 nM epinephrine at the conclusion of each trace distinguished α cells, which are robustly stimulated via β1 adrenergic receptors and β cells, which are potently inhibited via α2 adrenergic receptors. Intact imaging experiments were performed across 3 mice, with isolated islets analyzed independent of each other, before combined into the final dataset. Unless otherwise noted, a minimum of three mice were used for each experiment. Analysis was performed using whole α or β cell populations resulting in 100 to 1000 α or β cells per group.

### Immunofluorescence

Pancreases were isolated and cryosectioned as described previously [8]. For immunofluorescence of cryosectioned tissue sections, we followed a protocol identical to those previously published from our group [7, 8, 31]. Whole mount staining of intact islets following live imaging was carried out over a 5-day process as described previously [7, 8]. Slides and whole mount islets were imaged on a Nikon Eclipse Ti using a 60x lens with oil. Primary antibodies were validated in previous studies [7, 8, 32] from our lab or against tissue known to lack the antigen for the target antibody. All antibodies and concentrations used are available in supplemental table 1.

### Analysis and Quantification of SSTR3 Positive Cilia across islet cell lineages

Fixed Image analysis of membrane GFP or membrane tdTomato colocalization with SSTR3 was performed in ImageJ [33]. Each SSTR3 positive cilium was marked in ImageJ, then compared against both the GFP and tdTomato acquisitions of the same field to align cilia with respective fluorescence. Cilia with no overlapping signal were recorded as “unidentified.” All cilia imaging files names were scrambled prior analysis to ensure rigor.

### Analysis of live imaging results

Multi-image time series of islet behavior were acquired as previously described [7, 8, 34]. For analysis, Python v3.8 was used, (code available at https://github.com/Huising-Lab). To account for islet drift when imaging, an affine stabilization with a temporal window size of 10 frames was applied to every image of every time series. Each field of view in each imaging experiment was flattened into a median projection of the intensity. This projection was used to segment individual cells using CellPose [35]. Cells were segmented into regions of interest (code available at https://github.com/Huising-Lab), then eroded by 10% to ensure no overlap between individual cell regions of interest (ROIs) within the islet. Raw mean intensity values were then used to quantify ROI specific behaviors during live imaging, then normalized on a per ROI basis. Cell identity was assigned through a first filter using a T test for significance comparing the epinephrine response value against the 5 minutes immediately preceding epinephrine. After this filter, post-hoc immunofluorescence of glucagon and insulin confirmed cell specificity.

### Adenovirus Generation and Transduction

Recombinant jRGECO1a [36] (Addgene#100852) adenoviral particles were generated using the AdEasy system [37]. Adenoviral particles were delivered at a multiplicity of infection of 50 for a period of 12-16 hours to dissociated islet cells.

### Statistical Analysis

Comparisons made within the same slide sample for fixed image analysis or time series of the same cell were performed using a paired student’s T-test (p < 0.05). For comparisons made between populations of cells, the student’s T-test (p <0.05) was used. All analyses were performed in GraphPad Prism (Version 10) with raw data processed in Python (v3.8).

## Results

### Alpha cells express SSTR2 and beta cells express SSTR3 in mice

Previously published datasets from our lab have identified the specific SSTR mRNA profiles for FACS-purified α, and β cells of the islet (Fig.1a). We validated these results by immunofluorescence for SSTR2, which only colocalizes with glucagon positive α cells (Fig. 1b). Protein expression of SSTR3 was limited to the primary cilia (Fig. 1c). To assign cell identity to individual SSTR3-positive cilia, we crossed mice expressing α, β or δ cell specific Cre drivers with floxed membrane-TdTomato/membrane-GFP mice. This resulted in mice where the cell membrane including the cilia for each specific lineage was labeled with GFP, while the lineage-negative cilia expressed TdTomato (Fig. 1d). Quantification of the colocalization between these signals confirmed that most β cell cilia express SSTR3, However, to our surprise, no SSTR3 protein was detected on α cells and very little on delta cells (Fig. 1e).

**Figure 1:**
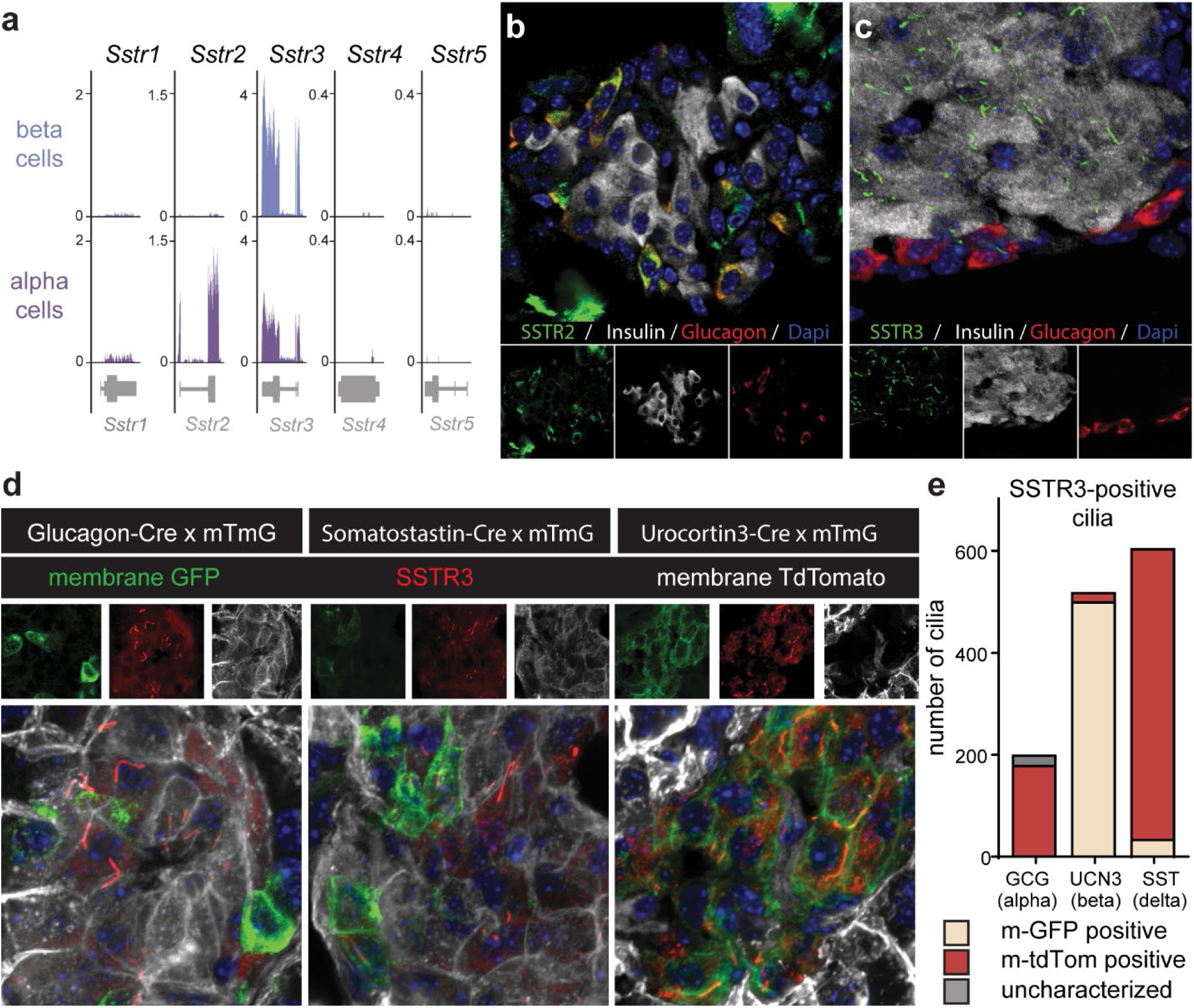
Alpha cells express SSTR2 and beta cells express SSTR3 in mice. Bulk α and β cell RNAseq results identify *Sstr3* message present on both α and β cells and *Sstr2* message present only in the α cells (a), with *Sstr1, Sstr4* and *Sstr5* message absent from both cell types. Immunostaining of SSTR2 and SSTR3 protein expression identifies SSTR2 on exclusively α cells (b), while SSTR3 is expressed on primary cilia of islet cells (c). SSTR3 positive cilia cell identity was confirmed against colocalization with a membrane GFP signal delivered by cell specific Cre driver. Of cilia observed, the vast majority of cilia that overlapped with the GFP cell identity marker were those from the β cell (d). The Membrane TD-tomato is pseudocolored white in panel D to accentuate the overlap between SSTR3 and the membrane GFP. Quantification of the overlapping SSTR3 and membrane GFP or membrane TD-Tomato confirms the majority of SSTR3 positive cilia are localized to the β cell of the islet (e).

### Somatostatin signaling attenuates Ca^2+^ in α cells more than in β cells

Given this difference in SSTR profiles between α and β cells, we compared the ability of SST to inhibit the Ca^2+^ response that was activated in both cell types simultaneously. We imaged islets isolated from constitutive GCaMP6s reporter mice with a progressive stimulation with 16.8 mM glucose and an amino acid mixture (AAM) (2 mmol/L of L-glutamine, L-alanine and L-arginine each) to reflect mixed-meal (carbohydrates and protein) post-prandial conditions, which tonically activates Ca^2+^ in both α and β cells (Fig. 2a). Subsequent application of SST (100 nM) caused a significant drop in the average Ca^2+^ signal intensity in both α and β cells (Fig. 2b,c) that was greater in α cells compared to β cells (Fig. 2d). Cell identity was validated by a combination of the response of each individual cell to epinephrine (which activates α and inhibits β cells) and post-hoc immunofluorescence for insulin and glucagon (Fig. 2e). Still images from this imaging experiment (Fig. 2f) are accompanied by a movie of a second islet from a separate mouse following the same experimental design (Movie S1).

**Figure 2:**
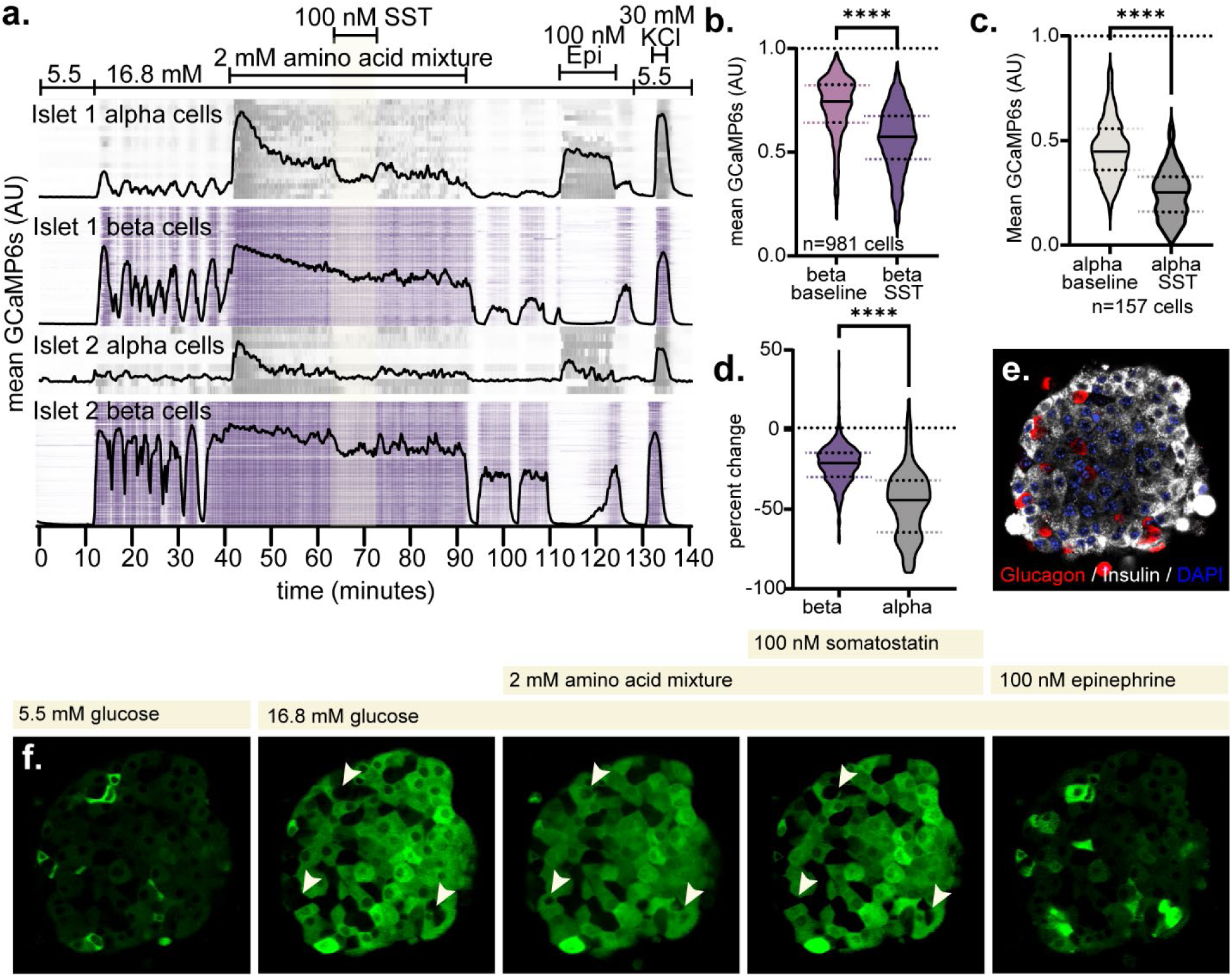
Somatostatin signaling attenuates Ca^2+^ in α cells more than in β cells. 17 beta-actin-Cre x GCaMP6s islets were analyzed across 3 separate mice, resulting in analysis of 157 α and 981 β cells. Islets were stimulated using both 16.8mM glucose and 2mM of an Amino Acid Mixture to drive Ca^2+^ transients. Two representative islet responses shown (a). A 10 minute treatment with 100 nM SST significantly reduced both α and β cell Ca^2+^ intensity compared to the 10 minute baseline intensity immediately preceding treatment (b,c) (Paired Student’s T test, Significance < 0.0001) Comparison of the percent drop in GCaMP6s intensity reveals a significantly larger effect of SST in inhibition of α cell Ca^2+^ (d) (Student’s T test, Significance < 0.0001). Representative immunofluorescence of imaged islet reveals α and β cell fate as stained by glucagon and insulin (e). Still images of representative islet highlight selectiveness of metabolic stimuli highlight cell behavior in response to experimental stimuli (f-j). White arrows (g-i) indicate an α identified by both epinephrine response and immunofluorescence. An additional islet is presented (g) with α and β cell specific ROIS isolated from each other to highlight cell behavior and fidelity. These cell specific ROIs are merged for ease of visualization of α and β cell behavior (h).

### Dissociation of intact islets increases β cell sensitivity to somatostatin

In the intact islet configuration, gap junction coupling between β cells could present an effective “resistor” to individual β cell Ca^2+^ inhibition by SST. In parallel, the cilia location of SSTR3 on β cells may function to compartmentalize downstream cytosolic Ca^2+^ [38, 39]. To differentiate between these scenarios and increase the scale of our imaging capabilities, we dissociated constitutive GCaMP6s islets and imaged them in a monolayer culture across several thousand β cells (n=2922) and several hundred α cells (n=301). This experimental design removes gap junction connections through physical separation, while retaining SSTR3 expression on primary cilia. The overall response (Fig. 3a) was comparable to their responses in the intact islet configuration (Fig. 2a). In both cell types, SST significantly inhibited Ca^2+^ in the dissociated state, with a more robust effect again observed in the α cells (Fig.3b-d) However, SST was a more robust inhibitor of Ca^2+^ in dissociated β cells (Fig. 3b) compared to β cells in the intact islet (Fig. 2a, 3e) as expected. Interestingly, Ca^2+^ responses in α cells, which are not connected via gap junctions to each other or to β cells, is also more robustly inhibited in dissociated α cells compared to intact (Fig. 3c, 3f). Still images from a representative experiment illustrate the effect of SST on α and β cells (Fig.3g-j) and confirm cell identity (Fig. 3j, k) (Movie S2).

**Figure 3:**
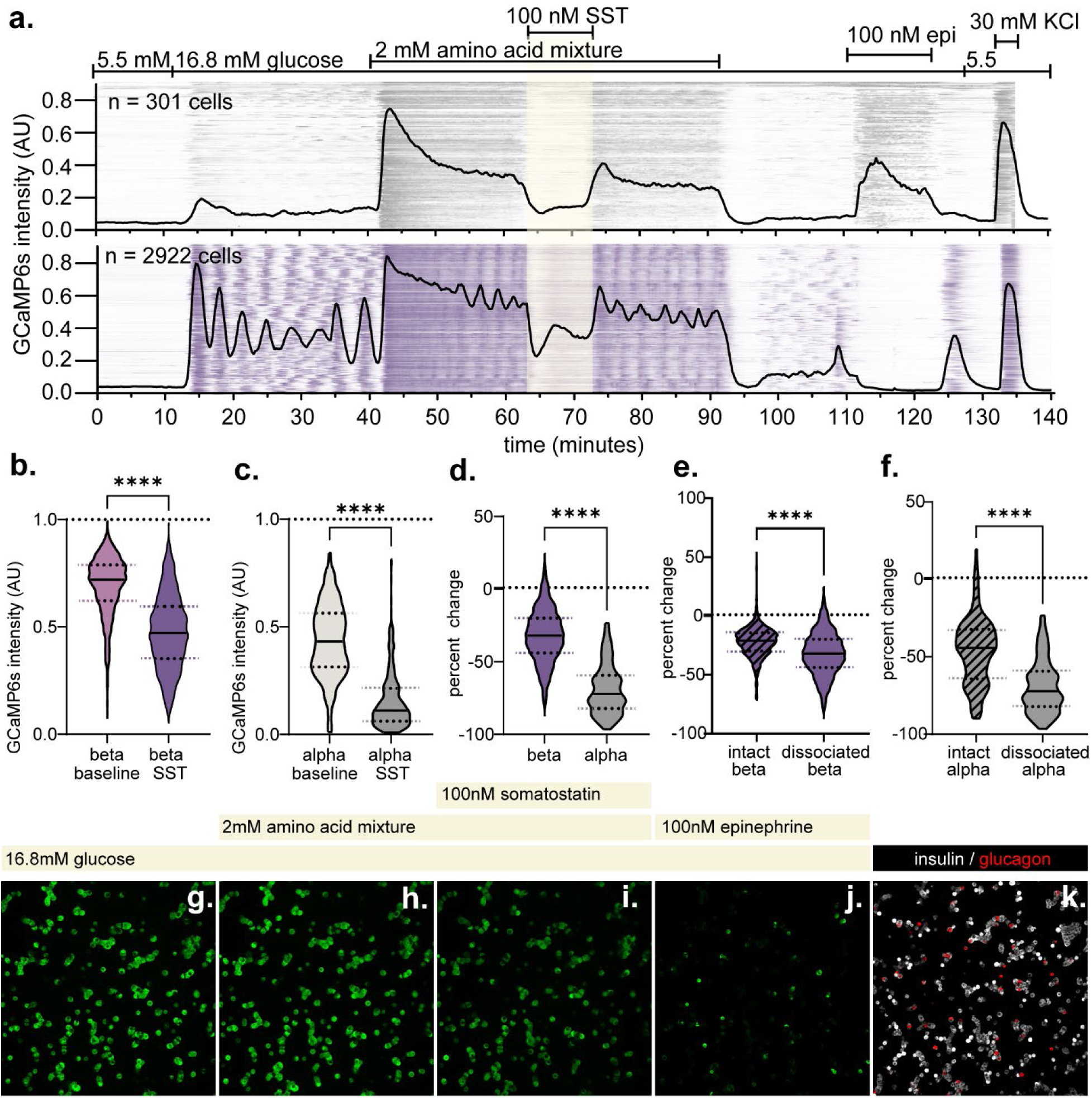
Dissociation of intact islets Increases β cell sensitivity to somatostatin. 3 separate dissociated cultures from pooled beta-actin-Cre x GCaMP6s mice were generated (10 mice total). In total, 2922 β cells and 301 α cells were assessed across imaging experiments. Representative Ca^2+^ intensity in response to stimuli identical to the intact islets is presented across one imaging experiment, with all cells combined in one graph and overlaid intensity plot (a). Treatment with 100 nM SST results in a significant drop in GCaMP6s fluorescence in both α and β cells when comparing the SST treatment period to the 10 minute baseline in the mixed meal treatment immediately preceding(b,c) (Paired Student’s T Test, <0.0001). A direct comparison of the percent change during SST treatment reveals a significantly larger drop in dissociated α cells compared to β (d) (Student’s T Test, <0.0001). Comparing the change in intact β and intact α against dissociated cell populations reveals both are more sensitive to 100 nM SST treatment when dissociated (e,f) (Student’s T Test, <0.0001) Still images (g-j) illustrate cell behavior in response to metabolic stimuli and treatments. Immunofluorescence of glucagon and insulin coupled with epinephrine response identifies α and β cells (j,k).

### Somatostatin robustly attenuates cAMP in β and α cells

Having determined that SST inhibited Ca^2+^ generated by robust nutrient stimulation with greater potency in α than β cells, particularly in intact islets (Fig. 2), we next determined the ability of SST to inhibit cAMP in both cell types. We stimulated islets from constitutive CAMPER (cCAMPER) reporter mice with gastric inhibitory polypeptide (GIP) to activate a robust cAMP response in both α and β cells, which each express the Gαs-coupled gastric inhibitory polypeptide receptor (GIPR) (Fig. 4a) [16, 40]. This allowed for quantification of the inhibition of cAMP upon addition of SST (100 nM) in ROIs for α cells (blue) and β cells (red) (Fig. 4b) (Movie S3). Somatostatin robustly inhibited cAMP levels to a similar extent in α and β cells (Fig. 4c,d). A combination of post-hoc immunofluorescence on cCAMPER islets (Fig. 4e,f) coupled with epinephrine response confirmed cell identity (Fig. 4b). Individual α and β cell behaviors are captured as pseudocolored cell specific ROIs in response to GIP, SST and Epinephrine (Fig. 4 g-j)

**Figure 4:**
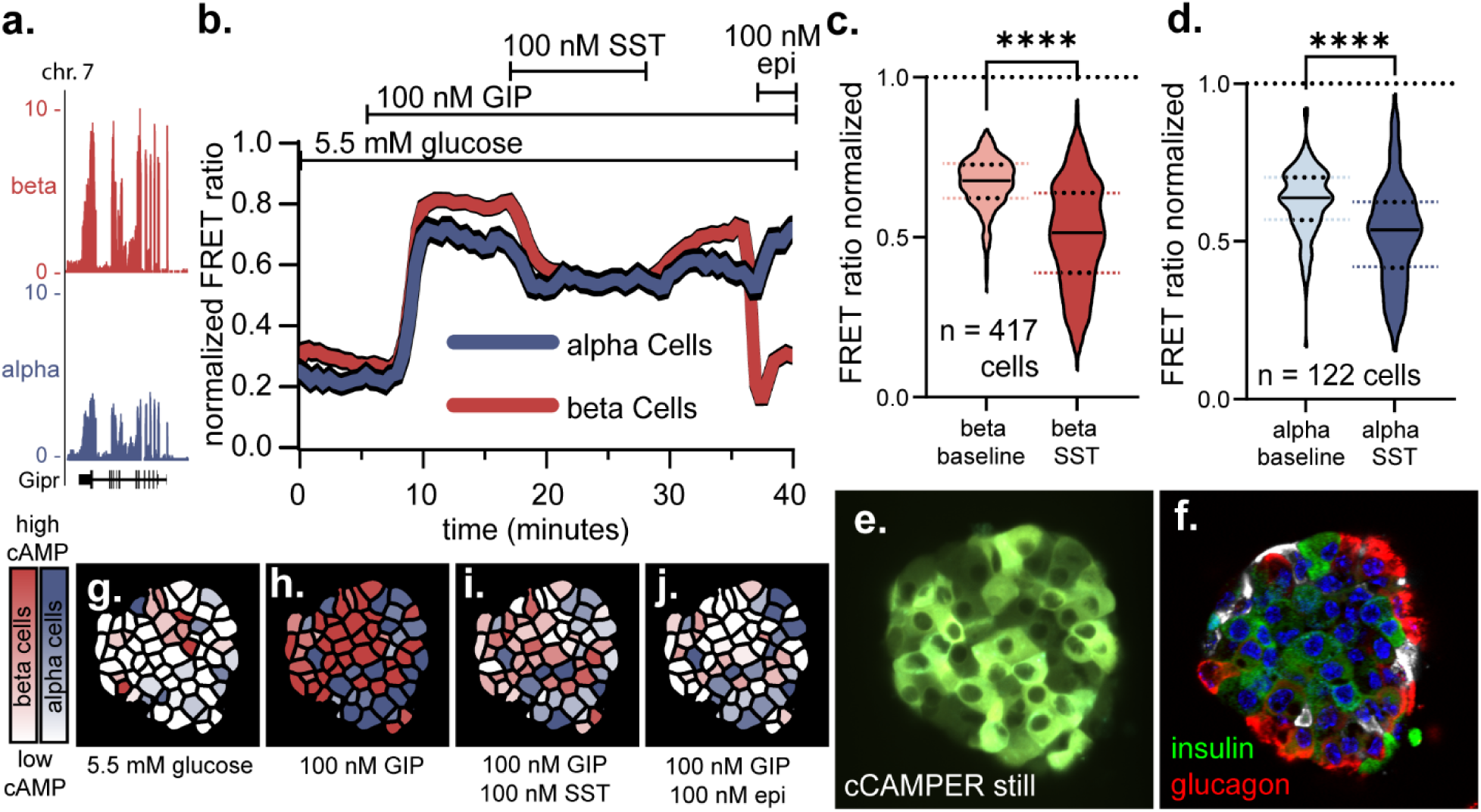
Somatostatin robustly attenuates cAMP in β and α cells. Mouse α and beta cell transcriptomes identify meaningful *gipr* expression in both cell types (a). 9 beta-actin-Cre x CAMPER islets were imaged from 3 mice on separate days. Across all imaging experiments, 417 β and 122 α cells were imaged. Example data from one islet is presented during treatment in basal glucose (5.5mM) with 100 nM GIP stimulating cAMP production in α and β cells, and 100 nM SST applied to track inhibition (b). Treatment with 100 nM SST resulted in a significant decrease in cAMP in both α and β cells (c,d) (paired student’s T Test, <0.0001). Representative still of cAMP expressing islet and corresponding immunofluorescence confirm cell identity and islet structure (e,f). The islet response to GIP and SST is presented as pseudo colored cell masks to track both cell types and changes over time (g-j).

### Selective SSTR inhibition relieves Ca^2+^ attenuation in α and β cells

To validate the receptor specificity of the inhibitory effects of SST on α (SSTR2) and β cell (SSTR3) Ca^2+^, islets from GCaMP6s reporter mice were stimulated with a mixed meal as before (Fig. 2). Stimulation with SST (100 nM) was applied to establish a new “inhibitory baseline” against the mixed meal stimulus (Fig. 5a). Application of the selective SSTR2 antagonist 406-028-15 (500 nM) [41] and the SSTR3-selective antagonist SST-ODN-8 (500 nM) [42] both significantly relieved α cell Ca^2+^ inhibition (Fig. 5b,c). Antagonism of SSTR2 also initially relieved β cell Ca^2+^ immediately followed by a sharp decrease in Ca^2+^ inhibition that is unique to β cell Ca^2+^ in response to SSTR2 antagonism (Fig. 5a). This led to a statistically significant overall reduction in β cell Ca^2+^ compared to the inhibitory baseline (Fig. 5d). Application of the SSTR3 antagonist SST-ODN-8 relieved SST-mediated Ca^2+^ inhibition in β cells as expected (Fig. 5e). Visualization of whole islet Ca^2+^ behavior in response to the mixed meal, SST, and antagonist captures the specificity of the antagonists in shaping α and β cell response to SST (Fig. 5f-l) (Movie S4). To highlight the effect of these treatments on individual cell response, cell specific ROIS were mapped to highlight both α (Fig. 5g) and β (Fig. 5h) cell behaviors and subsequently merged (Fig. 5i).

**Figure 5:**
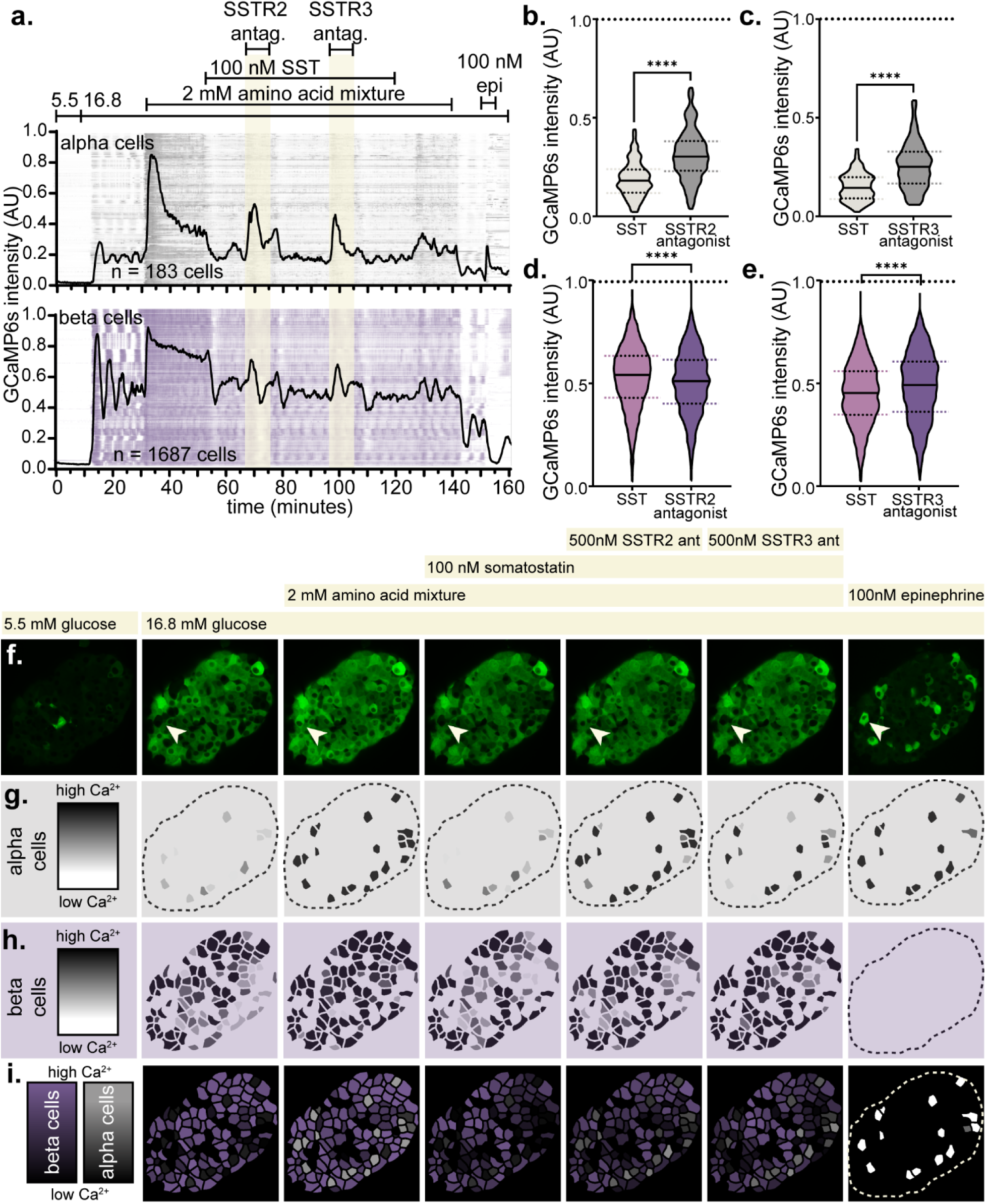
Selective SSTR inhibition relieves Ca^2+^ attenuation in α and β cells. 16 beta-actin-Cre x GCaMP6s islets from 3 separate mice were imaged, resulting in data acquired from 183 α and 1687 β cells. Islets were subjected to the same series of metabolic stimuli as in figure 1, but with a prolonged 100 nM SST treatment to establish baseline. The SST mediated baseline against the mixed meal background was used to introduce 500 nM of SSTR2 (406-028-15) and SSTR3 (SST-ODN-8) antagonists. Line graphs and intensity plots represent the average intensity of all cells acquired and each individual cell. (a). Both the SSTR2 and SSTR3 antagonists yielded significant relief of SST mediated Ca^2+^ inhibition in α cells (b,c) (paired student’s t-test, <0.0001). In Β Cells, SSTR2 antagonism resulted in a significant decrease in Ca^2+^ from the already suppressed SST baseline (d) (Paired Student’s T-Test, <0.0001). Treatment with the SSTR3 Antagonist significantly relieved SST attenuation of Ca^2+^ in β cells (e) (Paired Student’s T-Test, <0.0001). Still Images highlight a representative islet response to stimuli and peptides over the course of an imaging experiment (f). White arrow (f) highlights a representative α cell across numerous treatments. Cell specific ROIs were mapped to α and β cell specific ROIs to highlight cell behavior across treatments (g,h). Mapped ROIs were merged to ease in visualization of separate cell behaviors (i).

### Selective SSTR inhibition relieves cAMP attenuation in α and β cells

Next, we quantified the effects of SSTR2 and SSTR3 using selective inhibitors on α and β cell cAMP responses. Islets from cCAMPER mice were first stimulated with 100 nM GIP to increase cAMP in both α and β cells, followed by inhibition by 100 nM SST. Against this tonic SST inhibition, we applied consecutive pulses of the SSTR3 antagonist SST-ODN-8 and the SSTR2 antagonist 406-028-15 at 500 nM each (Fig. 6a). As expected, blocking SSTR3 relieved the inhibition of cAMP by SST on β cells (Fig. 6b). However, selective blockade of SSTR2 resulted in a significant reprieve from SST-mediated inhibition of cAMP in β cells (which do not express SSTR2) (Fig. 6c). In α cells, SSTR3 antagonism had no effect on SST mediated inhibition of cAMP (Fig. 6d), while the SSTR2 antagonist significantly relieved SST mediated inhibition of α cell cAMP as expected (Fig. 6e), in line with mRNA (Fig. 1a) and protein expression of SSTR2 (Fig. 1b). Pseudocolored islet maps generated from cell specific ROIs highlight the specificity of GIP, SST, the SSTR antagonists, and epinephrine in shaping α cell (blue) and β cell (red) cAMP response (Fig. 5f-k) (Movie S5).

**Figure 6:**
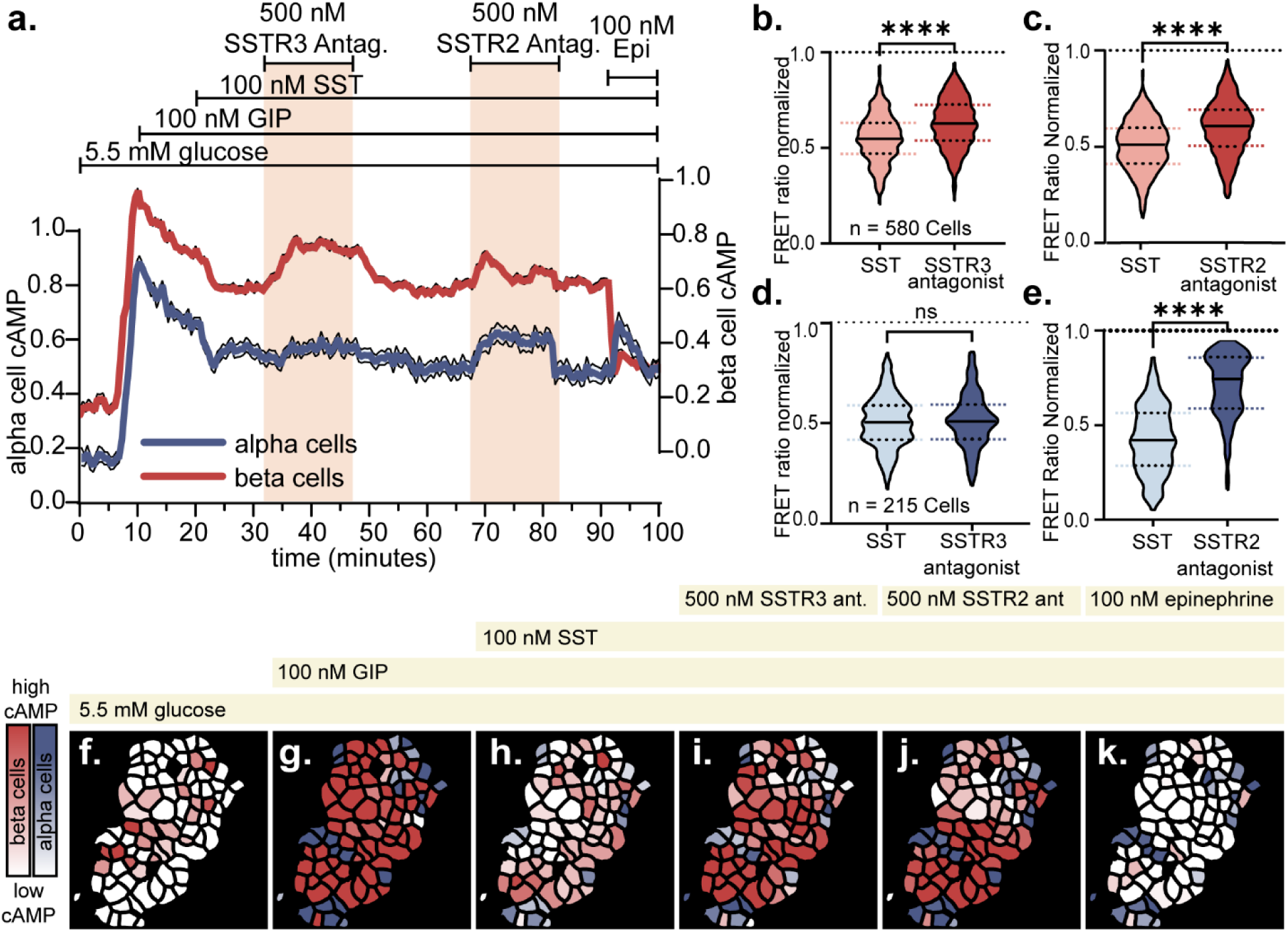
Selective SSTR inhibition relieves cAMP attenuation in α and β cells. 15 beta-actin-Cre x CAMPER islets were imaged from 4 separate mice with a total cell count of 215 α cells and 580 β cells. A tonic SST inhibition baseline was established against the background of basal glucose (5.5mM) and sustained GIP (100 nM). (a) Against this background, β cell cAMP was significantly increased in response to SSTR2 and SSTR3 antagonism (b,c) (paired student’s T test, <0.0001). In α cells, SSTR3 antagonism had no effect on cAMP, but SSTR2 antagonism significantly elevated cAMP activity (d,e) (paired students T test, <0.0001). Pseudo colored images of an islet treated with the stimuli in (a) are shown to highlight changes in α and β cell cAMP (f-k).

### Islet dissociation removes SSTR2-mediated relief of β cell cAMP

We hypothesized that dissociating and physically separating intact islet cells into a monoculture would dilute any paracrine contributions from α cell glucagon to β cell cAMP. For this experiment we used Ucn3-Cre x lsl-CAMPER mice, which exclusively report cAMP in β cells. We co-transduced these dissociated cultures with an adenovirus delivering jRGECO1a, a red shifted orthogonally fluorescent Ca^2+^ sensor to allow the simultaneous tracking of Ca^2+^ and cAMP in the same population of β cells (Fig. 7a). We again observed robust inhibition by SST of GIP-induced cAMP and glucose-induced Ca^2+^ (Fig. 7b,c). Application of the SSTR3 antagonist SST-ODN-8 significantly relieved SST-mediated inhibition of Ca^2+^ and cAMP in β cells (Fig 7d,e). However, there was no longer a significant change in β cell Ca^2+^ or cAMP following SSTR2 antagonism (Fig. 7f,g). Still images of these treatments with both Ca^2+^ and pseudocolored cAMP ROI maps are presented (Fig. 7h-o) (Movie S6).

**Figure 7:**
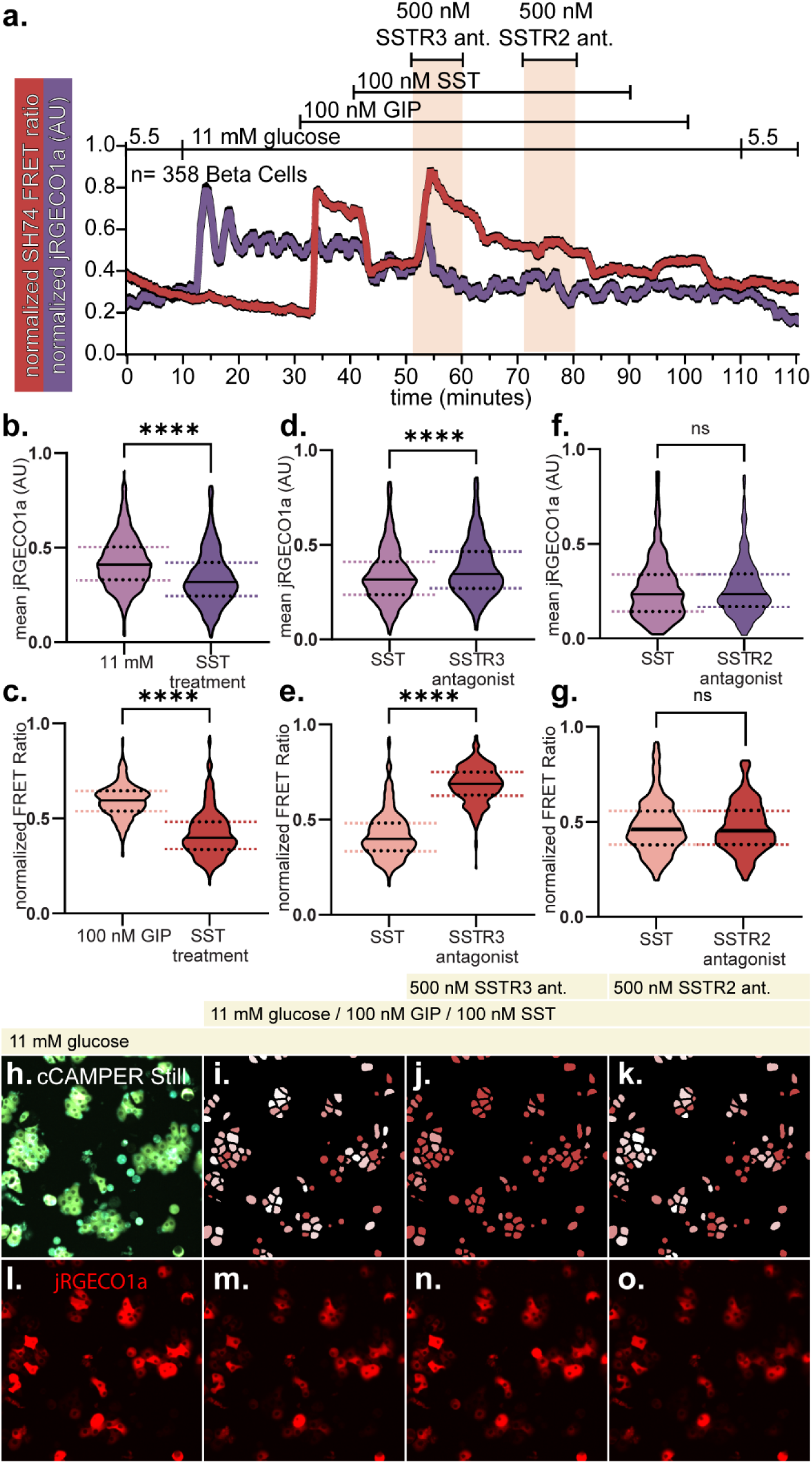
Islet dissociation removes SSTR2-mediated relief of β cell cAMP. To assess β cell Ca^2+^ and cAMP simultaneously, a β cell specific Cre driver was crossed with the CAMPER mouse and islets transduced with the red Ca^2+^ sensor jRGECO1a. This experiment was performed with pooled islets from two mice dissociated and seeded together. This yielded 358 total β cells. Islets were dissociated and subjected to the same strategy of exciting the cells and re-establishment of SST baseline to test SSTR2 and 3 antagonist effects (a). In dissociated β cells, Ca^2+^ and cAMP was significantly inhibited by 100 nM SST (b,c), with inhibition of both relieved by the SSTR3 antagonist (d,e)(Paired students’ T Test, <0.0001). Addition of the SSTR2 antagonist resulted in no significant relief of SST inhibition of either cAMP or Ca^2+^ (f,g) (paired students’ T Test). Representative dissociated cultures expressing the genetically knocked in CAMPER sensor and virally delivered jRGECO1a sensor are displayed (h-o).

### Blocking SST inhibition of α cells potentiates insulin release by glucagon and β cell GLP1R

To dissect our live imaging observations and place them in context with islet hormone secretion, we conducted static secretion experiments for glucagon and insulin from the same islets. Islets were treated with the same stepwise mixed meal stimuli used during Ca^2+^ imaging experiments. Stimulation with high glucose alone resulted in insulin secretion and caused a significant reduction in glucagon release, in line with prior observations [29] (Fig. 8a,b). The addition of amino acids (2 mM) on top of high glucose resulted in a modest, but statistically significant increase in glucagon release compared to high glucose alone (Fig. 8a), that did not affect insulin secretion. However, blocking α cell SSTR2 caused a robust increase in glucagon and insulin secretion (Fig. 8a,b). To determine if the large increase in glucagon was responsible for the insulinotropic effect of SSTR2 inhibition, we added Exendin9-39 to prevent α cell glucagon from activating β cell GLP1Rs. Doing so fully blocked insulin release, while glucagon secretion remained elevated (Fig. 8a,b). Exendin9-39 by itself did not alter glucagon or insulin secretion under mixed meal stimulation.

**Figure 8:**
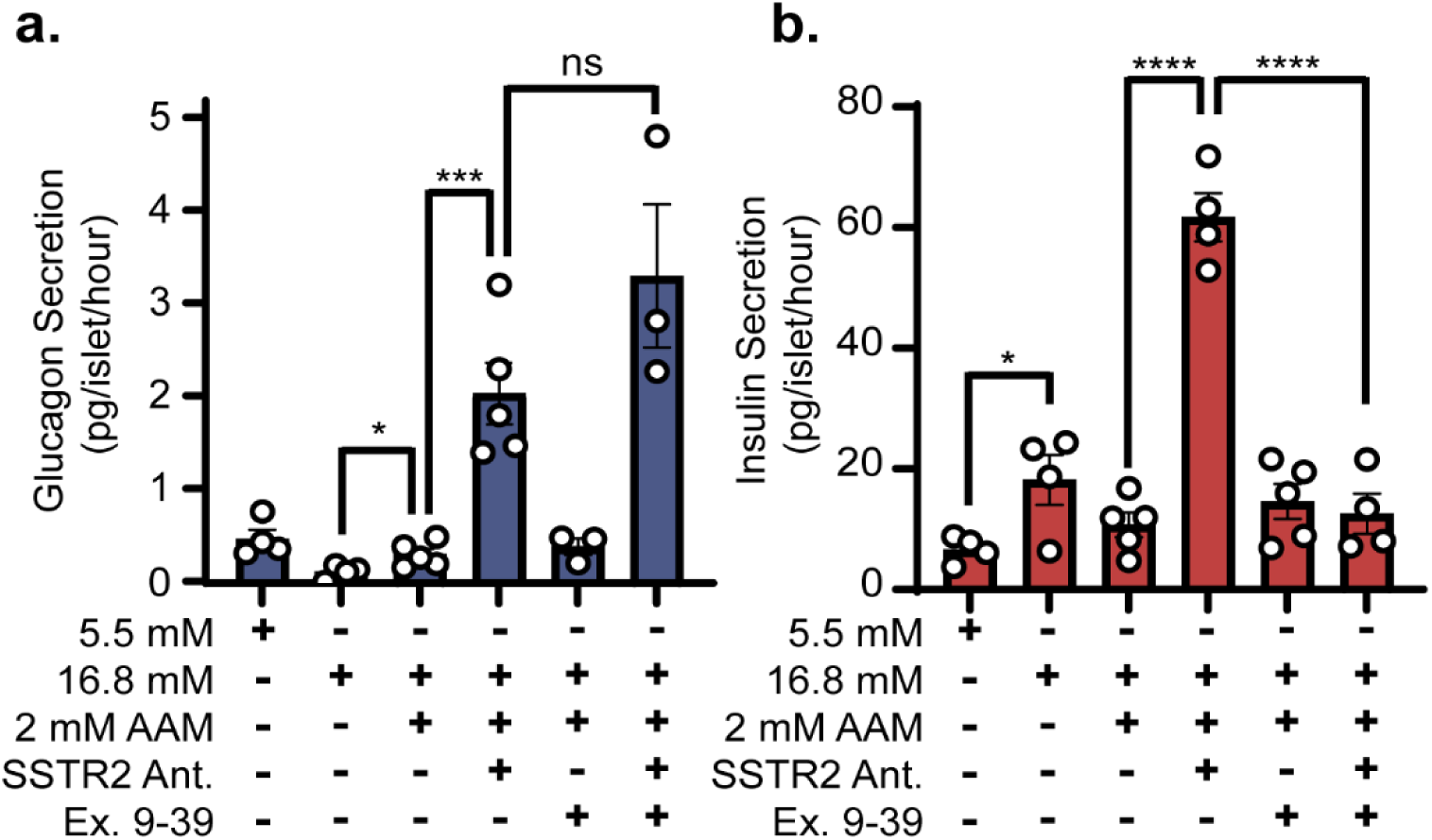
Blocking SST inhibition of α cells potentiates insulin release by glucagon via β cell GLP1R. Insulin and Glucagon secretion was acquired from a pooled islet population from 12 mice. Glucagon secretion was significantly increased when adding amino acids to a high glucose solution (a). Glucagon secretion was significantly higher in the presence of 500 nM SSTR2 antagonist (a). There was no significant difference in glucagon secretion with the addition of the GLP1r antagonist Exendin 9-39 (500 nM) (a). Insulin secretion was significantly increased between low (5.5mM) and high (16.8mM) glucose solutions. Insulin secretion was significantly higher in the presence of 500 nM SSTR2 antagonist (b). Addition of Exendin 9-39 resulted in a significant decrease in insulin secretion in the presence of the SSTR2 antagonist (b). All comparisons between groups were performed with the Student’s T test between samples. Significance was calculated as p < 0.05.

## Discussion

Within the last several years, our understanding of the α cells of the pancreatic islet has shifted from their original textbook definitions as counter regulatory cells responding to hypoglycemia [2], to a more cooperative relationship with β cells under nutrient stimulation [9, 43]. How this revised view of α cell biology intersects with δ-cell feedback during the post-prandial phase has remained unclear. We sought to understand how the inhibitory effect of the δ cell shapes individual α and β cell behavior as well as their paracrine relationship through somatostatin-mediated attenuation of cAMP and Ca^2+^. To accomplish this, we first validated cell specific transcriptome expression by visualizing the SSTR3 protein exclusively on β cell cilia despite robust *Sstr3* mRNA expression in all three islet cell types [16]. We confirmed expression of *Sstr2* mRNA and SSTR2 protein on α cell surface that are fully in line with our own and other groups’ prior observations [16–19].

### SSTR2 is a more potent inhibitor of cell activity than SSTR3

We assessed the relative ability of somatostatin to inhibit α and β cells and whether this depended on the inhibition of α and β cell Ca^2+^ and cAMP. Traditionally, most work in the islet field has focused on delivering genetically encoded sensors reporting on cAMP or Ca^2+^ to a single cell type using specific Cre drivers [44, 45]. To enable the simultaneous tracking of cAMP or Ca^2+^ across the full islet, we instead opted to express genetically encoded sensors for cAMP and Ca^2+^ in both α and β cells across the entire islet, and adopted work flows to be able to deconvolute the responses by each cell type with high fidelity [7, 29].

Simultaneous imaging of Ca²^+^ in both α and β cells revealed that SST produced a substantially larger decrease in α cell Ca²^+^ than in β cells. The muted β-cell Ca²^+^ response likely reflects two factors; first, SSTR3 on the β cell is restricted to the cilia, limiting its’ localized compartment of action [46, 47]. Second, once robustly activated by nutrient stimulation, gap junction connections between β cells of the intact islet may insulate these cells from the inhibitory effects of SST [7, 48]. Dissociation of islets removed this gap-junctional insulation and increased β cell SST sensitivity, affirming the hypothesis that gap junction coupling buffers individual β cells from SST. Notably, α cell Ca²^+^ remained more strongly inhibited than β cell Ca²^+^ even in dissociated cultures. This indicates an additional cell-autonomous feature that increases α cell SST sensitivity over β cell sensitivity which we predict is a result of differences between membrane bound SSTR2 and ciliary SSTR3 signaling.

In both islet cell types, cAMP increases driven by GIP were inhibited at a similar magnitude by SST. In contrast to its effect on Ca²^+^, the ciliary localization of SSTR3 therefore does not limit whole cell cAMP decreases in β cells, in line with recent results that documented the ability for ciliary GPCRs such as SSTR3 to readily modulate whole cell cAMP [38]. However, ciliary GPCRs are known to engage distinct downstream pathways including transcriptional programs compared to membrane-localized receptors such as SSTR2 [39, 49]. This difference in downstream signaling was beyond the scope of this study, and presents another mechanism through which SST mediated inhibition of α cells via cell surface SSTR2 and β cells via ciliary SSTR3 may differ.

### SST inhibition of prandial insulin release requires alpha cell-mediated paracrine feedback

Application of an SSTR3 antagonist selectively blocked the SST effect on β cell cAMP and Ca^2+^. The SSTR2 antagonist blocked the SST effect on α cell Ca^2+^ and cAMP as expected. However, the SSTR2 antagonist also caused an increase in β cell cAMP, measured in intact islets. We hypothesized this increase in β cell cAMP was a result of the increase in local α cell glucagon acting on β cell GLP1Rs as recently documented [11, 50]. Dissociation of islets into single cell monocultures physically separated the α and β cells and abolished the increase in β cell cAMP following the inhibition of α cell SSTR2. These observations are in line with the presence of a paracrine factor, presumably glucagon, being released from the α cell under metabolic or incretin co-stimulation. To demonstrate that δ cell inhibition of α cells under prandial condition restricted downstream β cell insulin secretion we conducted static secretion experiments using intact islets. Application of SSTR2 antagonist to islets under mixed-meal stimulation resulted in a significant increase in glucagon release, in line with previously published results from islets isolated from whole body SSTR2 knockout islets [51]. However, application of an SSTR2 antagonist also resulted in a significant increase in insulin release, even though beta cells do not express SSTR2. Blocking of the GLP1 Receptor using exendin 9-39 did not affect glucagon secretion, but prevented the increase in insulin release during SSTR2 antagonist treatment. These data corroborate that the robust increase in β cell cAMP from intact islets following the inhibition of α cell SSTR2 was mediated via α cell glucagon activating β cell GLP1R. These data affirm that a central role of the α cell in the islet environment is the paracrine potentiation of insulin secretion from the neighboring β cells, as has been shown recently with both advanced imaging [43], and islet perifusion experiments [52]. Our data extend this understanding to demonstrate that under mixed-meal stimulation, δ cells achieve their attenuation of insulin secretion primarily via the indirect inhibition of this local paracrine α cell-dependent potentiation loop instead of via the direct inhibition of β cells via ciliary SSTR3. Lack of δ cell inhibition of this α to β paracrine potentiation of insulin secretion likely at least partially explains the excessive nutrient-stimulated insulin secretion observed following the deletion of δ cells or SST [31, 53, 54].

### Delta cell control under non-fasting and post-prandial insulin secretion

This work is distinct from our previously published observations which demonstrated that the non-fasting setpoint for glucose in-between meals is determined via direct and modest feedback inhibition by δ cells that is independent of α cells and right-shifts the β cell glucose threshold by 1 to 1.5 mM glucose [8, 31]. Future studies will and should expand on the composition of the post-prandial stimulation and the effect of incretin co-stimulation and how this recruits an α cell-dependent potentiation of insulin secretion that is under inhibitory control by δ cells.

### Limitations of the Study

We used SST in this study at a concentration approximately 10 times higher than its published IC50 for both SSTR2 and 3 [55] and adopted antagonist use at a 5 times higher molar ratio, as we have successfully in prior studies [31]. Our rationale for doing so was to achieve maximal levels of inhibition and to accommodate for possible loss of efficacy because the peptide would have to diffuse into the islet interstitial space given that our experiments were conducted on *ex-vivo* isolated islets that lack circulation.

The advantage of the SSTR2 and 3 specific antagonists that we used is their amenability to the live imaging approaches employed by us. We have used these published compounds in past publications [31]. The use of α and β cell specific SSTR2 and SSTR3 knockout islets would also be used as a companion approach for our observation. There are global knockout mice for both SSTR2 [51] and SSTR3 [56], but these constitutive whole-body null mice are potentially confounded by off target effects and the possibility of compensation from any of the other 4 SSTRs. If floxed *Sstr2* and *Sstr3* alleles became available, they could form a useful complement to the results we described here.

Future studies will also leverage the approaches here in human islets. Previous results examining electrical activity in human α and β cells identified SSTR2 as the functionally dominant receptor [57], which aligns well with our findings in mouse islets. However, the expression profile of SSTRs in human islets is much more heterogenous than mice [58]. As such, disentangling the individual contributions from the human islet SSTR network would be a substantial and separate undertaking.

### Conclusions

Through the examination of the effect of SST on the inhibition of cAMP and Ca^2+^ in α and β cells of mouse islets, we have quantified how α and β cell behaviors are attenuated at the secondary messenger level. By a combination of pharmacologically perturbing this system via selective antagonists and dissociation of islets to prevent paracrine interaction, we revealed that the largest contribution of SST on β cell insulin secretion under mixed-meal post-prandial conditions is achieved by the inhibition of the paracrine relationship whereby α cells potentiate nutrient-stimulated insulin secretion from β cells. Importantly, this is distinct from our previously published observations which demonstrated that the non-fasting setpoint for glucose *<u>in-between</u>* meals is determined via direct and modest feedback inhibition by δ cells that is independent of α cells and right-shifts the β cell glucose threshold by 1 to 1.5 mM glucose [8, 31]. Taken together, our findings serve to clarify our understanding of the complex direct and indirect paracrine relationships between the α, β and δ cell. While δ cells under non-fasting conditions control the glucose setpoint via modest direct inhibition of β cells [8], under post-prandial conditions δ cell feedback inhibition of insulin release is achieved predominantly via the inhibition of α cell-mediated potentiation of the β cell.

## Data availability

All data will be shared by the corresponding author upon reasonable request.

## Code availability

All code used for analyses and data processing are available on githhub (https://github.com/Huising-Lab).

## Contribution Statement

RGH and MOH designed research; RGH, JJL, KZ, SL, RC, AH ADN and MOH performed research; RGH, JJL, AH, and MOH analyzed data; RGH and MOH wrote and edited the manuscript. All authors approved of the final version of the manuscript. MOH is responsible for the integrity of the work as a whole.

## Disclosure of Authors’ relationships and activities

M.O.H. received funding from ThermoFisher Scientific for work unrelated to the results described in this manuscript.

## List of Abbreviations

Ca^2+^: Intracellular calcium
cAMP: cyclic adenosine monophosphate
SST: Somatostatin
GLP-1R: Glucagon-Like Peptide-1 receptor
SSTR2: Somatostatin Receptor 2
SSTR3: Somatostatin Receptor 3
α: “Alpha” cell
β: “Beta” cell
δ: “Delta” cell
GSIS: Glucose-stimulated insulin secretion
GPCR: G Protein Coupled Receptor
ROI: Region of interest
GIPR: gastric inhibitory polypeptide receptor
GIP: gastric inhibitory polypeptide

