## Supplemental Table 1 for "Somatostatin Receptors Shape Insulin and Glucagon Output within the Pancreatic Islet through Direct and Paracrine Effects"

| Antibody | Species | Concentration | Company | Catalog # |
|---|---|---|---|---|
| polyclonal anti-insulin | Guinea Pig | 1:500 | Dako | A0564 |
| monoclonal anti-insulin | Rat | 1:500 | R&D Systems Catalog | MAB1417 |
| Polyclonal anti-glucagon | Guinea Pig | 1:1000 | Progen | 16032 |
| Polyclonal anti Somatostatin | Sheep | 1:1000 | American Research Products | 13-2366 |
| polyclonal anti somatostatin receptor 2 | Rabbit | 1:1000 | Alomone | ASR-006 |
| polyclonal anti somatostatin receptor 3 | Rabbit | 1:1000 | ThermoFisher | PA3-207 |
| Polyclonal anti-GFP | Goat | 1:1000 | Rockland | 600-101-215 |
| Cy3-AffiniPure F(ab')2 Fragment Anti-Guinea Pig IgG | Donkey | 1:600 | Jackson Immunoresearch | 706-166-148 |
| 647-AffiniPure F(ab')2 Fragment Anti-Guinea Pig IgG | Donkey | 1:600 | Jackson Immunoresearch | 706-166-148 |
| Cy3-AffiniPure F(ab')2 Fragment Anti-Rat IgG | Donkey | 1:600 | Jackson Immunoresearch | 712-166-153 |
| 647-AffiniPure F(ab')2 Fragment Anti-Rat IgG | Donkey | 1:600 | Jackson Immunoresearch | 712-606-153 |
| 488-AffiniPure F(ab')2 Fragment Anti-Goat IgG | Donkey | 1:600 | Jackson Immunoresearch | 705-546-147 |
| 647-AffiniPure F(ab')2 Fragment Anti-Sheep IgG | Donkey | 1:600 | Jackson Immunoresearch | 711-606-152 |
| Cy3-AffiniPure F(ab')2 Fragment Anti-Rabbit IgG | Donkey | 1:600 | Jackson Immunoresearch | 711-166-152 |
| 647-AffiniPure Anti-Rabbit IgG | Donkey | 1:600 | Jackson Immunoresearch | 711-605-152 |
